# Binding Ability Prediction between Spike Protein and Human ACE2 Reveals the Adaptive Strategy of SARS-CoV-2 in Humans

**DOI:** 10.1101/2020.06.25.170704

**Authors:** Xia Xue, Jianxiang Shi, Hongen Xu, Yaping Qin, Zengguang Yang, Shuaisheng Feng, Danhua Liu, Liguo Jian, Linlin Hua, Yaohe Wang, Qi Zhang, Xueyong Huang, Xiaoju Zhang, Xinxin Li, Chunguang Chen, Jiancheng Guo, Wenxue Tang, Jianbo Liu

## Abstract

SARS-CoV-2 (severe acute respiratory syndrome coronavirus 2) is a novel coronavirus causing an outbreak of COVID-19 globally in the past six months. A relatively higher divergence on the spike protein of SASR-CoV-2 enables it to transmit across species efficiently. We particularly believe that the adaptive mutations of the receptor-binding domain (RBD) of spike protein in SARS-CoV-2 might be essential to its high transmissibility among humans. Thus here we collected 2,142 high-quality genome sequences of SARS-CoV-2 from 160 regions in over 50 countries and reconstructed their phylogeny, and also analyzed the interaction between the polymorphisms of spike protein and human ACE2 (hACE2). Phylogenetic analysis of SARS-CoV-2 and coronavirus in other hosts show SARS-CoV-2 is highly possible originated from Bat-CoV (RaTG13) found in horseshoe bat and a recombination event may occur on the spike protein of Pangolin-CoV to imbue it the ability to infect humans. Moreover, compared to the S gene of SARS-CoV-2, it is more conserved in the direct-binding sites of RBD and we noticed that spike protein of SARS-CoV-2 may under a consensus evolution to adapt to human hosts better. 3,860 amino acid mutations in spike protein RBD (T333-C525) of SARS-CoV-2 were simulated and their stability and affinity binding to hACE2 (S19-D615) were calculated. Our analysis indicates SARS-CoV-2 could infect humans from different populations with no preference, and a higher divergence in the spike protein of SARS-CoV-2 at the early stage of this pandemic may be a good indicator that could show the pathway of SARS-CoV-2 transmitting from the natural reservoir to human beings.

## Introduction

Coronavirus is commonly found in nature and infects only mammals and birds^1-3^, among 46 species, only seven of them are human-susceptible^4,5^. Aside from SARS-CoV and MERS-CoV that cause deadly pneumonia in humans by crossing the species barrier^3,6,7^, SARS-CoV-2 is now bringing a global pandemic of respiratory disease (COVID-19) within six months after the first case has been confirmed in Wuhan city of China^8^. It has been identified as a novel coronavirus that is a member of β-coronavirus in the family Coronaviridae, which is a positive single-stranded RNA virus with a protein envelope^2^. Up to date, COVID-19 caused by SARS-CoV-2 results in more than six million people infected and over 380 thousand deaths worldwide. Compared with SARS-CoV and MERS-CoV, SARS-CoV-2 spreads more rapidly and being highly infectious to humans^8-10^. It is crucial to understand the origin of this coronavirus and its strategy in adapting to human hosts so efficiently, moreover, to apply this knowledge for controlling this pandemic and developing effective therapeutics and vaccines against COVID-19.

By reconstructing phylogenomic relationships among various coronavirus^11^, it showed around 70% genome sequence similarity with SARS-CoV^11^, and more closely related to bat coronavirus RaTG13 in the spike (S) gene^1^. Given closely related to SARS-like coronaviruses, the genome structure of SARS-CoV-2 is similar to other beta-coronaviruses, which composed in order with 5’-replicase ORF1ab-S-envelope(E)-membrane(M)-N-3’ with numbers of open reading frames (ORFs) function resembles those of SARS-CoV^12^. It is noted that RaTG13 is highly similar to SARS-CoV-2 especially between genes, while they differed in some crucial genomic structures, one of the most notable features is that a polybasic (furin) cleavage site insertion (PRRA residue) at the junction between two subunits (S1, S2) of S protein ^1,2,12,13^. Although some studies show bats could be the reservoir host for many coronaviruses including SARS-CoV, the reservoir host of SARS-CoV-2 remains unclear ^4,10,14^. Given the global spread of this epidemic, it draws a lot of attention to reveal the origins of the pandemic event. The evolutionary of SARS-CoV-2 may explain its infectiousness and transmissibility among different animal hosts and provide evidence about whether this virus is natural or artificial.

Spike glycoprotein on the surface of SARS-CoV-2 is the key to enter the target cells, which forms homotrimers protruding from the surface to recognize host cell receptor and cause membrane fusion ^15^. Spike protein contains S1 and S2 subunits and the receptor-binding domain (RBD) exists on S1 which can bind to the peptidase domain (PD) of angiotensin-converting enzyme 2 (ACE2), while S2 is responsible for membrane fusion during viral infection^14,16^. In SARS-CoV, RBD in spike protein is the most diverse part of the whole genome, of which six amino acid (Y442, L472, N479, D480, T487, and Y4911) have been found to play a key role in binding to ACE2 receptor and further in transmissibility across species boundary^16^. Similar to SARS-CoV, RBD mutations acquired during adaption to different host cells along transmission have also been observed in SARS-CoV-2. Thus, dissection of the key mutations in spike protein RBD that affect binding to the ACE2 receptor would be important for understanding the molecular mechanism of how SARS-CoV-2 infect human cells. Recent studies show the ACE2 receptor in host cells would mediate the entry ability of SARS-CoV-2 by interacting with spike protein, and its binding capacity to spike protein of SARS-CoV-2 determines the transmissibility of this coronavirus across species particularly among humans ^14,17^.

The present study intends to 1) reveal the phylogenetic relationship of SARS-CoV-2 identified in the different population at the genomic level, and provide genetic evidence based on the structural protein-coding genes (S, M, N, E) for identifying the nucleotide variations of SARS-CoV-2 collected from different regions amid COVID-19 pandemic; 2) provide a prediction for revealing the adaptive mutations on spike protein of SARS-CoV-2 based on the specific variation of S gene, and finding the differences of stability of spike protein mutants and their affinities with the human ACE2 receptors; 3) explore whether there are variants of ACE2 in different populations which may affect the infectivity of COVID-19. To this end, we analyzed the variants of *ACE2* in a large cohort including 1000 Chinese local people and other human populations and identified polymorphisms that may influent on the binding between ACE2 and spike protein of coronaviruses, furthermore, we predicted the affinities of spike protein to binding ACE2 variants to understand whether those changes would render individuals resistant or susceptible to SARS-CoV-2 at the molecular level. Herein our study could explain part of the origin of SARS-CoV-2 at a phylogenetic level based both on whole genome and multiple key genes, to allow the elucidation of population risk profiles and also help advance therapeutics such as a rationally designed soluble ACE2 receptor for the management of COVID-19.

## Materials and Methods

2,147 genome sequences of coronavirus and its S, M, N, E gene sequence collected from 160 regions (50 countries) globally were downloaded from GenBank (https://www.ncbi.nlm.nih.gov/genbank/) and GISAID (https://www.gisaid.org/) databases and further filtered according to their completeness. The genome sequencing of a local donor has been completed in a laboratory from the local CDC, and the relevant data is stored in the local CDC laboratory database for further analysis and research. In this study, all the virus sequences were aligned with Clustal Omega (V1.2.3)^18^ and filtered by sequences containing continuous 15 Ns, and 327 non-repetitive sequences of S gene and 469 of S, N, M, E combined sequences of SARS-CoV-2 were extracted from the filtered genome sequences respectively. Iqtree (v1.6.12)^19^ was used to determine the most suitable model for each sequence dataset. We chose the maximum likelihood tree reconstructed using MEGA v5.2^20^. The parameter we performed with 1000 Maximum number of iterations and approximate Bayes test. Figtree v1.4^21^ was performed on editing and screening the evolutionary tree topology. The calculation of genetic distance (whole genome and individual genes) was carried out by Kimura two factor correction method for nucleic acid level calculation. To avoid the prediction error caused by the selection of outgroups with a far evolutionary relationship, the complex outgroup was adopted in this study, and the sequence of MERS-CoV and SARS-CoV were selected as outgroups to predict the genetic relationship.

*In silico* mutagenesis of SARS-CoV-2 spike protein receptor-binding domain bound to the ACE2 receptor complex (PDBID: 6M0J) was used to predict the influence of binding affinity and protein stability. The proposed residue sites were substituted to 19 other amino acids and an ensemble of the conformations (the number of conformations was limit to 25) was generated for each mutant by low-mode MD (Molecular Dynamics). MM/GBVI was applied to calculate the binding affinity of each conformation and ACE2 molecules. The force field used for calculation was OPLS-AA, and the implicit solvent was the reaction field (R-Field) model. All calculations were performed in MOE (Molecular Operating Environment) package^22^. The structure of ACE2 receptor (PDBID: 6M17) was used to perform *in silico* mutagenesis. The genomic variants in the human *ACE2* gene for different populations were downloaded from the gnomAD database (https://gnomad.broadinstitute.org/) The proposed residue sites were substituted to the amino acids that have the reported point mutations according to gnomAD. The parameters to run the prediction of polymorphisms in hACE2 binding to spike protein of SARS-CoV-2 performed as same as the previous one about the prediction on variants of spike protein binding to hACE2.

## Results

### Phylogeny of SARS-CoV-2

The Maximum Likelihood (ML) phylogenic trees were constructed based on 2,147 genome sequences of SARS-CoV-2 with SARS-CoV and MERS-CoV were selected as the out-groups (Figure 1). After alignment, we merged identical sequences into one clade with labels kept as one. The genomic tree showed all SARS-CoV-2 was closely related to the SARS-like virus found in the horseshoe bat from Yunnan (RaTG13), and a coronavirus from pangolin collected in Guangdong province was the sister taxon with RaTG13 which is closer to the virus from bat than to other pangolins from Guangxi. SARS-CoV-2 is not related closely with SARS-CoV at the nucleic level even they have over 72% sequence similarity. Moreover, SARS-CoV-2 strains from different regions were hard to solved completely based on genome sequences with multiple polytomies presented on the genome tree, particularly in the later time of this pandemic. On the contrary, the strains that closer to the original direction and towards the root position on the tree have higher bootstrap values on the branch and solved better, which may indicate that there is a consensus evolution as SARS-CoV-2 adapting to human hosts environment in the spread of COVID-19. The full phylogeny trees were found in supplementary Figures 1-3.

**Figure 1.**
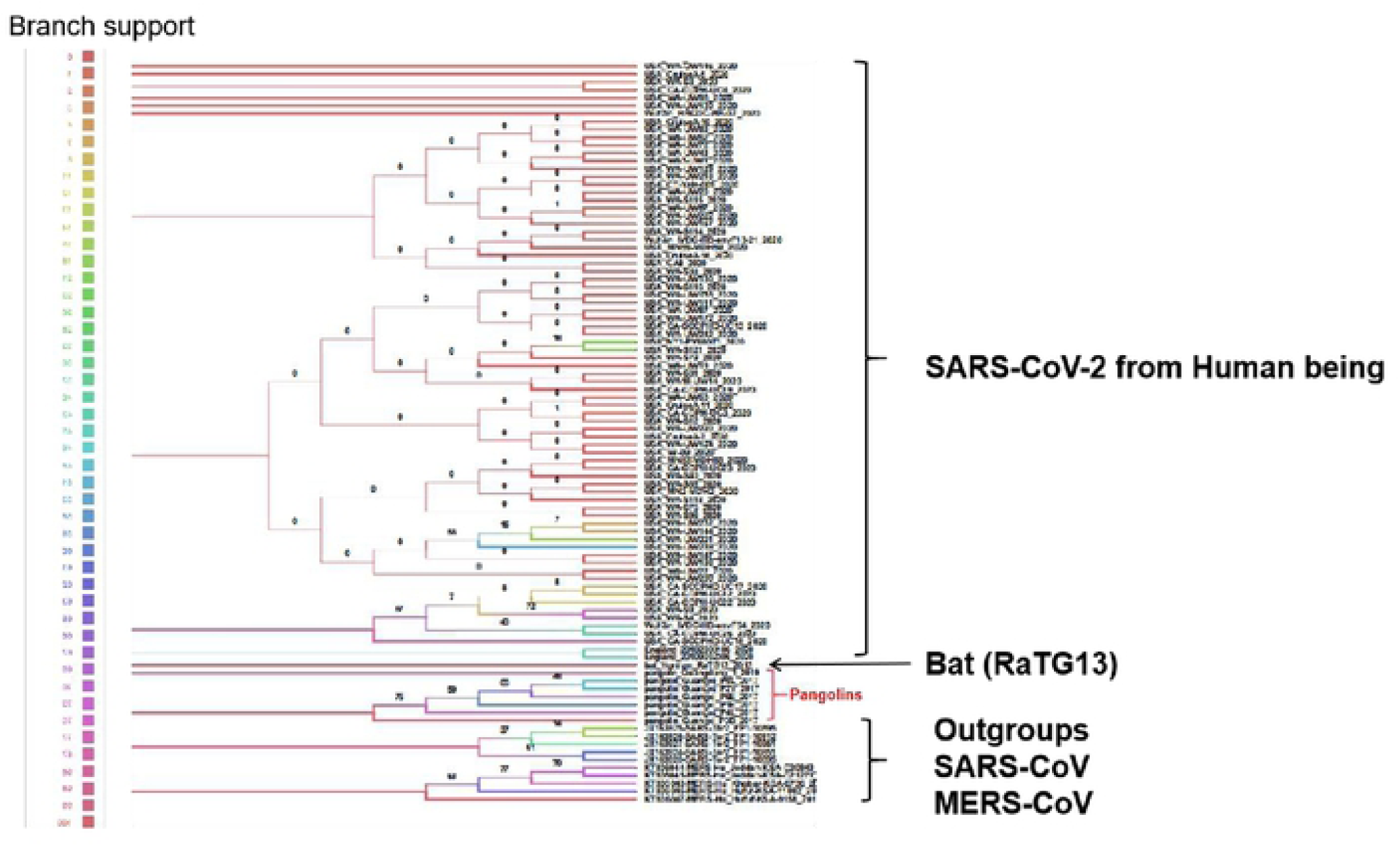
The ML phylogeny tree of different strains SARS-CoV-2 from various region all around the world (Partial, the full tree was found in Supplementary Figure 1), the bootstrap values were mapped on the branch as long as the colors annotated for all clades.

Besides, we reconstructed the phylogenetic trees based on multiple genes (S, N, E, M) (Figure 2), since structural protein-coding genes are fundamental to host cells recognition and infection. The phylogenetic relationship of multiple genes of different SARS-CoV-2 strains represented differently with the phylogeny based on the genome sequences, which might show the more adaptive pattern of the functional proteins rather than the evolutionary position of different strains from different host populations. Unlike the genomic phylogeny pattern, structural protein-coding genes showed more divergent on two polar areas than the strains in the middle part of the tree, we found a strain isolated from a local patient (HN-07) shared a high similar (> 98%) sequence with the ones from Australia, European countries and USA, and the case document showed this patient was confirmed after his seven-day trip to Italy and he didn’t show any symptoms before his journey. Some branches were poorly supported by the bootstrap values and exhibited polytomies, which indicated SARS-CoV-2 was adapting the human hosts globally and it is hard to determine the origin of SARS-CoV-2 solely based on nucleic data. Furthermore, same to the genomic tree, multiple genes sequences of SARS-CoV-2 are divergent from outgroups and we detected some mutations according to S gene sequences comparison (Figure 3).

**Figure 2.**
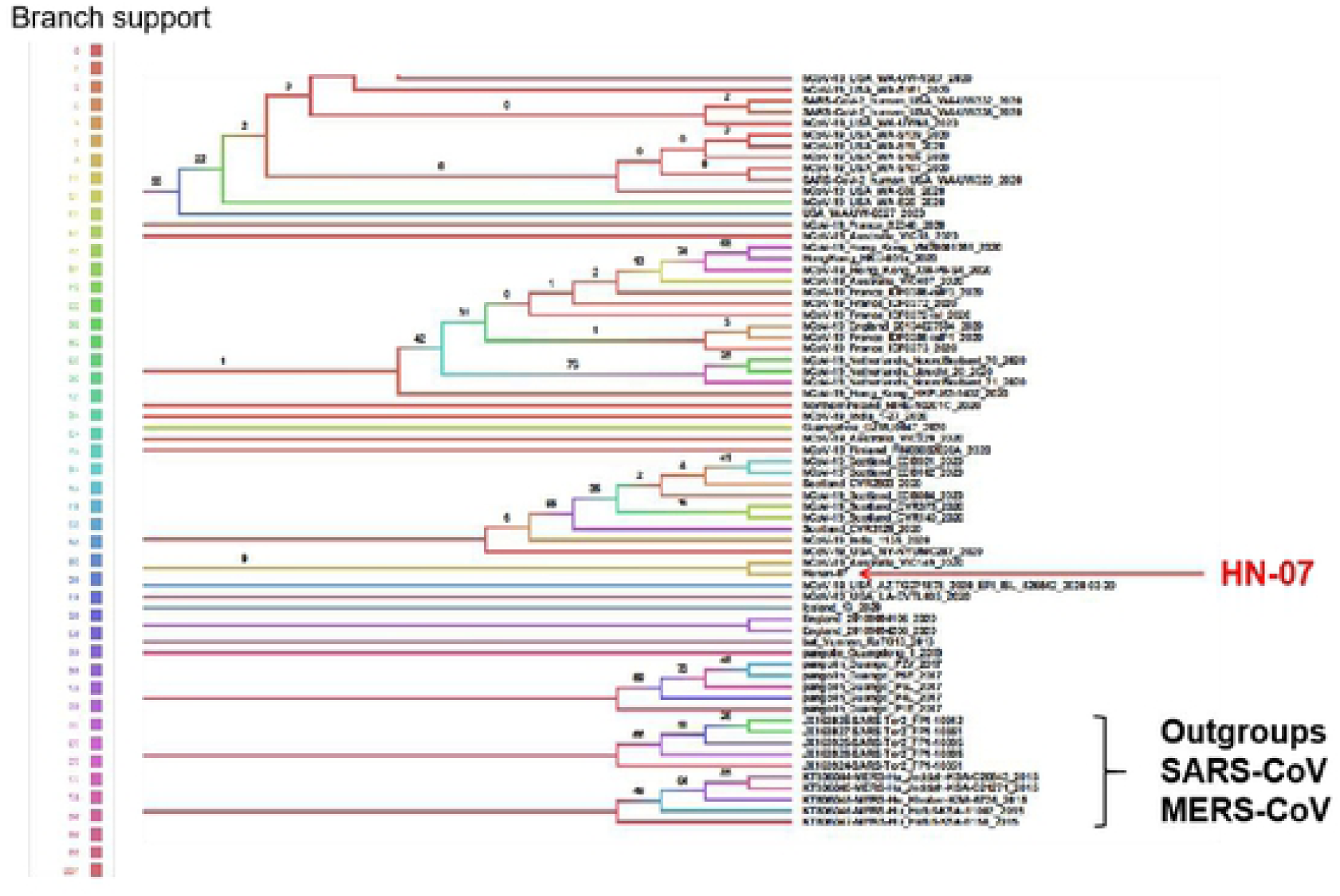
The ML phylogeny tree of S, N, M, N gene sequences of SARS-CoV-2 from various strains (Partial, the full tree was found in Supplementary Figure 2), the bootstrap values were mapped on the branch as long as the colors annotated for all clades.

**Figure 3.**
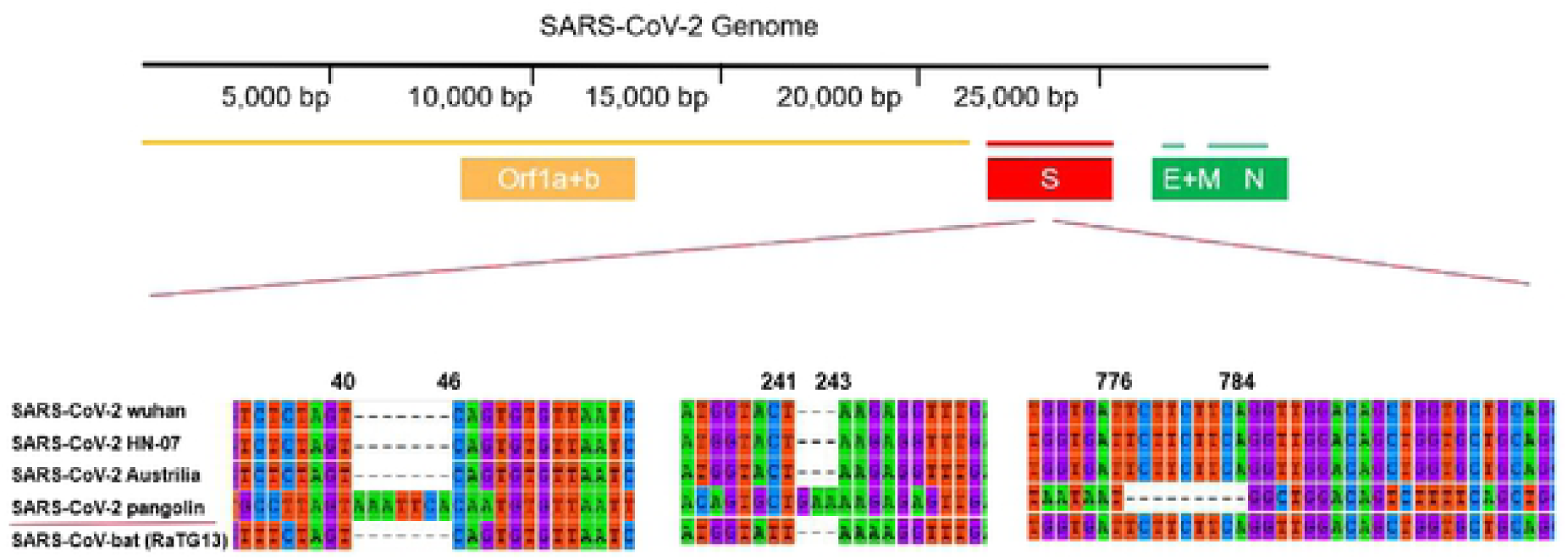
The alignment of S gene from different SARS virus.

As shown in Figure 3, we extracted and aligned the S gene sequences from SARS coronavirus in different hosts, the one from pangolin that closer to SARS-CoV-2 and RaTG13 was from a horseshoe bat (*Rhinolophus spp*) and we found no big fragment shift in S genes of pangolin and it was less similar to S in SARS-CoV-2 compared with RaTG13, but the insertions exhibited in pangolin indicated potential recombination in spike protein of coronavirus would occur during its cross-species hosts manumission.

According to S gene sequences in different strains, we found the similarity between SARS-CoV-2 and SARS-CoV-bat (98%) is higher than it compared with coronavirus from pangolins (85%). It showed more solved of the phylogenetic tree based on S gene (Fig. 4) compared to the trees based on genomic sequences and multiple genes. Nevertheless, the phylogeny reconstructed based on genome, S gene, and multiple genes, all indicate the SARS-CoV-2 is closely related to RaTG13 and the virus isolated from pangolin.

**Figure 4.**
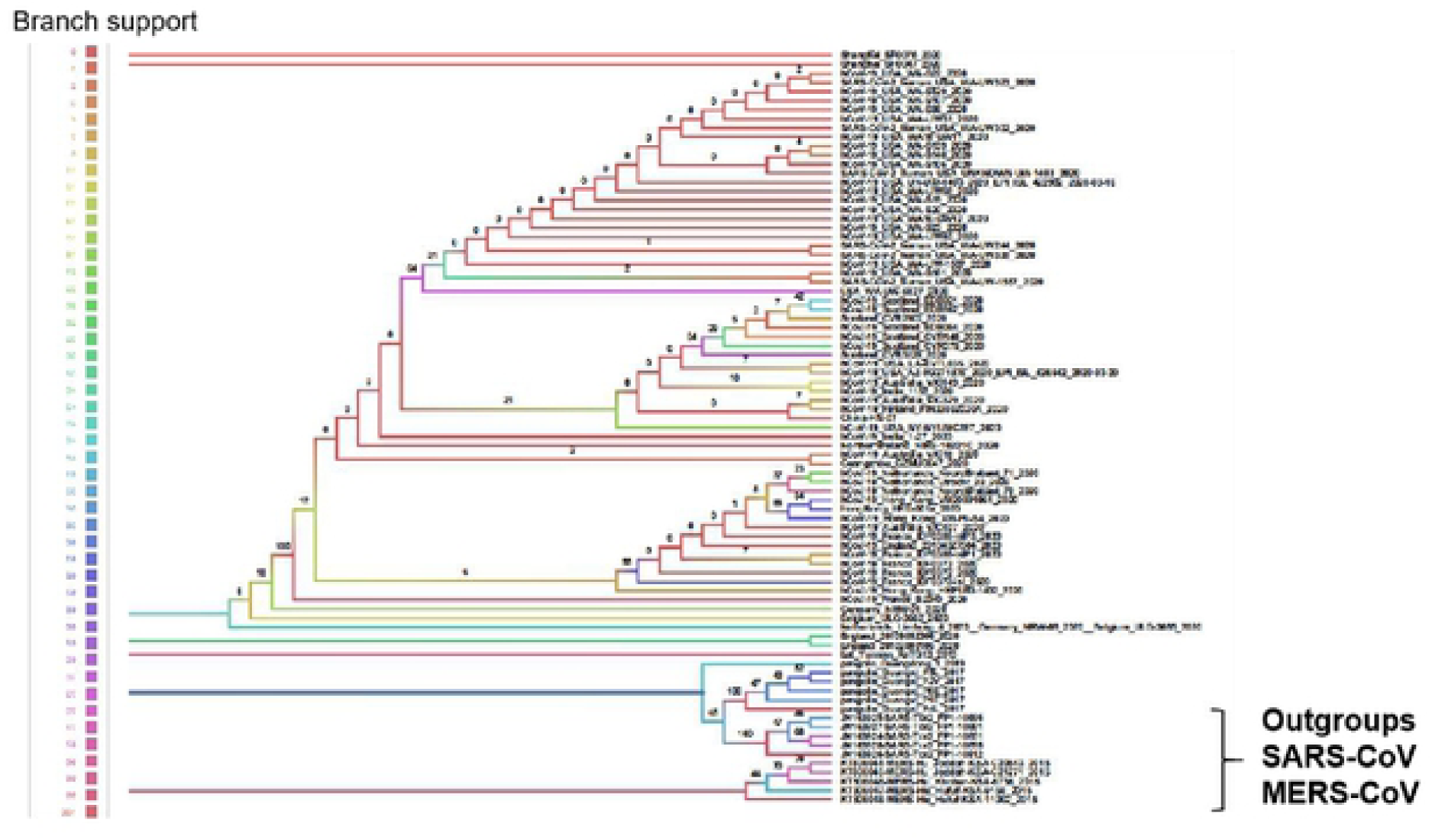
Phylogenetic tree based on S gene (Partial, the full tree was presented in Supplementary Figure 3), the bootstrap values were mapped on the branch as well as the colors annotated for all clades.

### Polymorphism prediction of spike protein in SARS-CoV-2

S gene has been studied as the key gene for SARS-CoV-2 binding to host cells’ receptor, and this gene shows less conserved compared to the genome sequence of SARS-CoV-2, and some studies suggest that the high divergence found in spike protein RBD and specifically the direct bonding sites to ACE2 receptor play an important role to SARS-CoV-2 adapting to different animal hosts or the populations in different regions. Combined with the phylogeny based on the S gene, we also detected all mutationson the S gene and their binding capacity with ACE2 receptor in human beings. 23 point mutations were predicted to significantly influence the affinity and stability of the spike protein (Fig. 5), among which 9 polymorphisms exhibited increased affinity and stability while 14 ones showed decreased affinity and stability (Table 1). All analysis outcomes of over 3000 polymorphisms in spike protein were shown in Supplementary Table 1.

**Table 1.** The affinity and stability of different variants of spike protein RBD bound to hACE2.

| **Mutants** | **Affinity**<br>**(kcal/mol)** | **dAffinity**<br>**(kcal/mol)** | **Stability**<br>**(kcal/mol)** | **dStability**<br>**(kcal/mol)** |
| --- | --- | --- | --- | --- |
| K417W | -90.17 | -3.68 | -3096.01 | 1.08 |
| G446W | -98.31 | -7.56 | -3100.18 | -0.42 |
| Y449A | -88.71 | 5.75 | -3098.95 | 2.41 |
| Y453Q | -85.63 | 3.36 | -3100.47 | 1.40 |
| L455A | -82.79 | 3.01 | -3101.17 | 2.30 |
| L455M | -92.15 | -6.35 | -3102.42 | 1.05 |
| F456A | -80.25 | 4.66 | -3100.11 | 3.05 |
| F456G | -80.11 | 4.80 | -3099.85 | 3.31 |
| A475W | -101.02 | -7.18 | -3101.49 | 0.28 |
| F486A | -84.93 | 6.49 | -3099.87 | 1.99 |
| F486G | -84.72 | 6.70 | -3099.50 | 2.37 |
| N487H | -86.01 | 5.65 | -3098.24 | 0.79 |
| Y489W | -96.85 | -5.74 | -3101.93 | 0.35 |
| Q493A | -90.20 | 3.63 | -3099.00 | 1.21 |
| Q493G | -89.33 | 4.50 | -3098.68 | 1.52 |
| G496A | -85.87 | 4.21 | -3102.19 | -0.18 |
| Q498A | -83.68 | 3.42 | -3100.48 | 1.45 |
| Q498W | -92.51 | -5.41 | -3102.17 | -0.25 |
| T500W | -96.40 | -5.19 | -3101.31 | 0.27 |
| N501G | -85.81 | 3.33 | -3098.31 | 1.94 |
| N501Y | -92.70 | -3.55 | -3100.42 | -0.17 |
| G502Y | -95.82 | -6.55 | -3102.78 | -0.84 |
| Y505A | -82.39 | 5.33 | -3101.12 | 1.98 |

**Figure 5.**
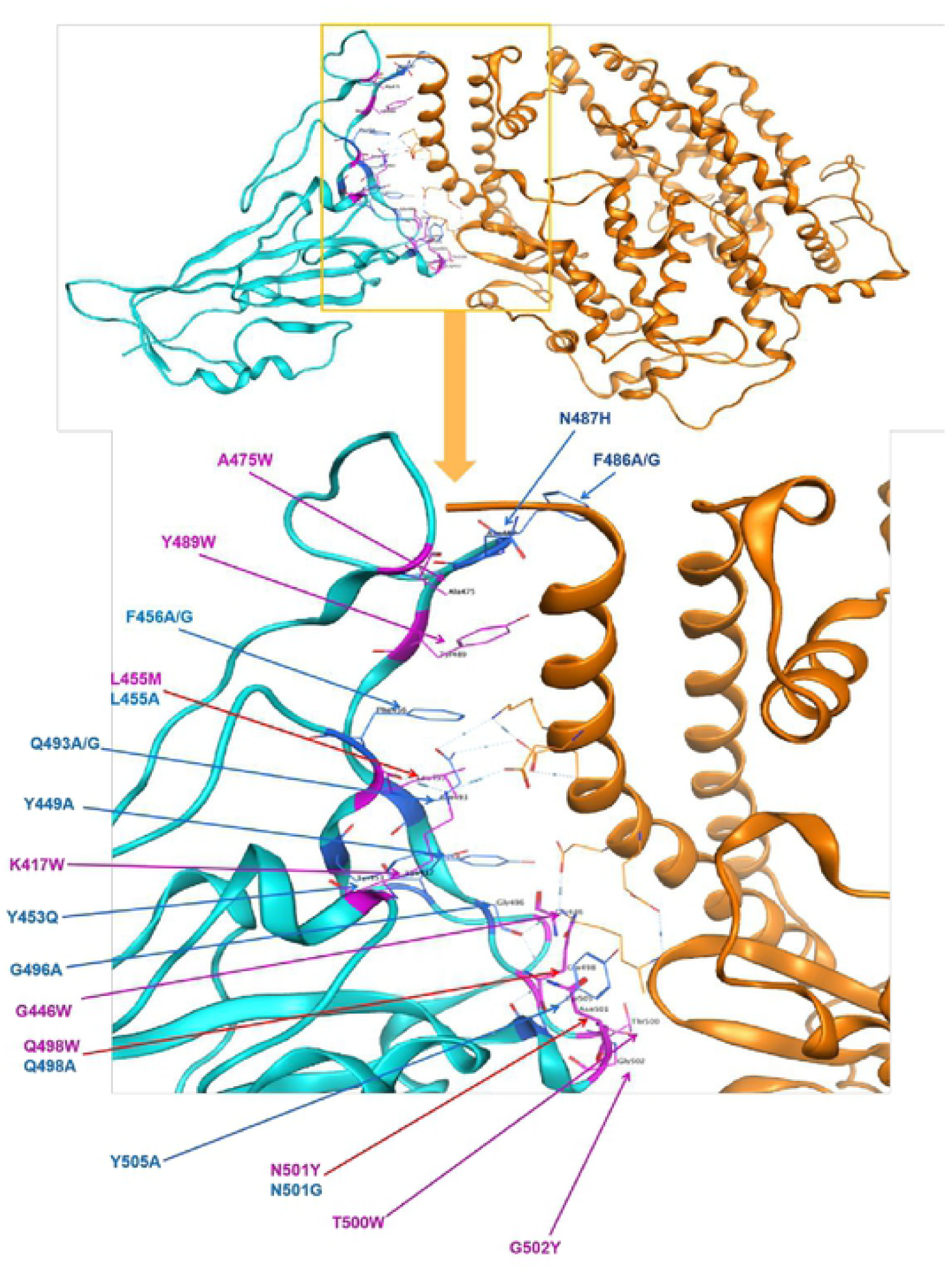
Identified polymorphism in spike protein RBD mapped to the structure of spike protein in SARS-CoV-2 in complex with ACE2 in humans. Cyan = spike protein, Orange=ACE2 direct bounding to spike protein. 23 point mutations causing affinity significant change on direct bounding of spike protein of SARS-CoV-2, blue presents decreasing affinity while purple shows increasing.

Figure 5 and Table 1 showed the missense mutation in spike protein RBD that has significant changes (Cutoff =3) of the affinity and stability with ACE2 receptors. Interestingly, residues L455, Q498 and N501 would have two potential mutations that leading to contrary affinities, including increased affinity in leucine changed to methionine on AA455, glutaurine changed to tryptophan on AA498 and asparagine changed to tyrosine on AA501, reduced affinity in leucine changed to alanine, glutamine changed to alanine, asparagine changed to glycine. The polymorphisms on the same residue causing two opposite effects of affinity suggested that mutations on those positions may lead to different adaptation directions on SARS-COV-2 to fit in different hosts. Furthermore, Phe456, Gln493, and Phe486 showed two mutations that both result in affinity reduction. The stability of spike protein with different mutations was showed in Table 1, we found only 5 mutants increased the stability of spike protein including G446W, G496A, Q498W, N501Y and G502Y, which is not consistent with the affinity.

<u>T</u>o match the mutations of spike protein that reported by SARS-CoV-2 database of China national center for bioinformation (https://bigd.big.ac.cn/ncov) with our predictions, a total of 1,150 polymorphisms in spike protein were collected from the database and 643 missense variants were selected and run analysis about their affinity and stability binding to ACE2 (Supplementary Table 2). We focused on 76 missense variants in region T333-C525 of spike protein and 9 variants bring significant changes on the affinity of it to ACE2 (Figure 6A), moreover, 13 variants cause structural stability changes in spike protein RBD (Figure 6B).

**Figure 6.**
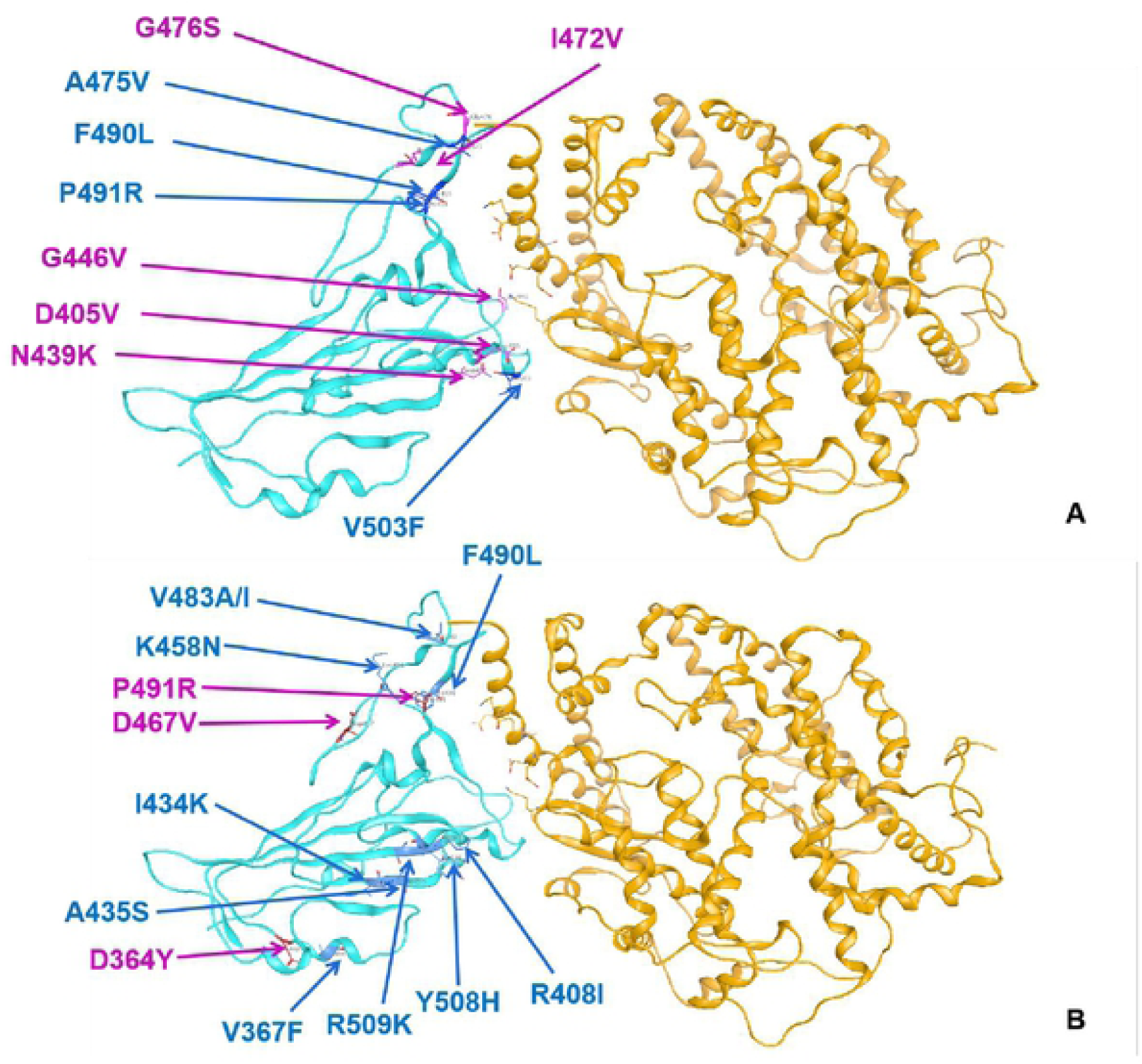
Reported polymorphism in spike protein RBD mapped to the structure of spike protein in SARS-CoV-2 in complex with ACE2 in humans. Cyan = spike protein, Orange=ACE2 and blue presents decreasing affinity and stability while red shows increasing ones. A: Affinity; B: Stability.

### Polymorphisms in ACE2 affect the binding ability to spike protein of SARS-CoV-2

We calculated the population frequency of 388 missense variants in ACE2 collected from GnomAD (Supplementary Table 3) and the local population (Table 2). Analysis of their stability and affinity to spike protein of SARS-COV-2 were performed in this study, and the results showed no significant differences in both affinity and stability.

**Table 2.** Analysis of affinity and stability of polymorphism in ACE2 from local population binding to spike protein of SARS-COV-2.

| Mutants | Affinity<br>(kcal/mol) | dAffinity<br>(kcal/mol) | Stability<br>(kcal/mol) | dStability<br>(kcal/mol) |
| --- | --- | --- | --- | --- |
| I468V | -65.09 | 6.88E-12 | -3107.75 | 0.85 |
| N638S | -65.09 | -4.76E-12 | -5930.30 | 0.33 |
| R708Q | -64.99 | 0 | -2906.26 | 2.17 |

## Discussion and Conclusions

Revealing the evolutionary origin of SARS-CoV-2 is of great value to understand its transmission pathway cross-species and provide a guide to long-term infection prevention of zoonotic coronavirus. Since it is a novel coronavirus, the extremely limited morphological information of SARS-CoV-2 could be used in the phylogenic analysis, genomic data of SARS-CoV-2 nowadays provide a helpful approach to identify its divergence along with the transmission among humans ^1,4^. The present study collected around 2,147 genome sequences of SARS-CoV-2 from over 50 countries and reconstructed their phylogeny based on the genome, S gene, and S-N-E-M gene sequences, it is consistent about the relationship between SARS-CoV-2 and the outgroup (SARS-CoV and MERS-Cov) which they are more closely related to SARS-CoV compared with MERS-CoV. The genome of SARS-CoV-2 is about ∼80% similar to SARS-CoV^11^ and ∼66% similar to MERS-CoV with low query coverage as 34%, but due to the structural and biological differences among them, they are strikingly different species of coronaviruses^9,23^. Some studies show the stability and viability of SARS-CoV-2 are higher than SARS-CoV generally in both aerosol environments and objects surfaces^24^, which indicates SARS-CoV-2 might adapt to human transmission better while MERS and SARS-CoV failed to do so^6,9,23,25^. Same with other studies^1,2,11,26^, we found all SARS-CoV-2 in our analysis were sister taxon with a coronavirus from horseshoe bat (RaTG13) which highly suspected as the natural reservoir host of coronavirus ^17,26^ and the similarity between SARS-CoV-2 and SARS-bat is high as 96.12%. Meanwhile, the coronavirus isolated from pangolins also were detected 91.02% identical with SARS-CoV-2 at the nucleotide level, which makes people think pangolins might be a potential intermediate host between the natural reservoir and human beings. However, according to the alignment of S gene sequence from different hosts, there are multiple insertions and deletions found in SARS-pangolin compared with S genes from SARS-CoV-2 and SARS-bat, which suggests the pangolin might not be the direct or only intermediate host and also explained by the low branch supports on phylogenetic trees. Unique furin (polybasic) cleavage site insertion (PRRA) were found at the in-between region of S1 and S2 subunits of spike protein of SARS-CoV-2, and this structure has potentials to enhance the infectivity of SARS-CoV-2, furthermore, this insertion has not been identified in SARS-pangolin and SARS-bat which also may indicate the essential intermediate host of SARS-CoV-2 remains unidentified ^14,15,27^.

Spike protein is a key at the first step of the viral infection in host cells for receptor recognition to most SARS-coronavirus^5,17^, and the S gene is more divergent in coronavirus at a genetic level relative to other genes^14^. Along adapting to the human host cellular environment, mutations occur in spike protein in some of which enable to enhance the binding affinity and stability of spike protein-hACE2 complex, and the enhancement would increase the transmissibility of SARS-CoV-2 among human and bring more severe disease^16^. For instance, the transmissibility and pathogenicity of SARS-CoV are lower from strains isolated in 2003-2004 compared with the ones from 2002-2003^28,29^, and MERS just faded away literally by itself along infecting human beings^9^. We performed affinity and stability of spike protein RBD (T333-C525) of SARS-CoV-2 by using MOE2019 (Molecular Operating Environment) to human ACE2 (S19-D615), point mutations on spike protein were simulated with amino acid scanning and 3860 mutations were set up for analysis (Supplementary Table 1). 23 variants of spike protein RBD that causing significantly higher or lower affinity to ACE2 and stability were focused in this study, the cutoff we set up was three for determining significance (Table 1), among 9 of the mutations that highly enhance the binding between spike protein and ACE2, the residue G446 (Figure 7) was also reported CDC up to date (Figure 6, Supplementary Table 2), and this sample documented as collected in early March with closer to the original direction of the phylogenetic tree based on genomic data (figure 1).

**Figure 7.**
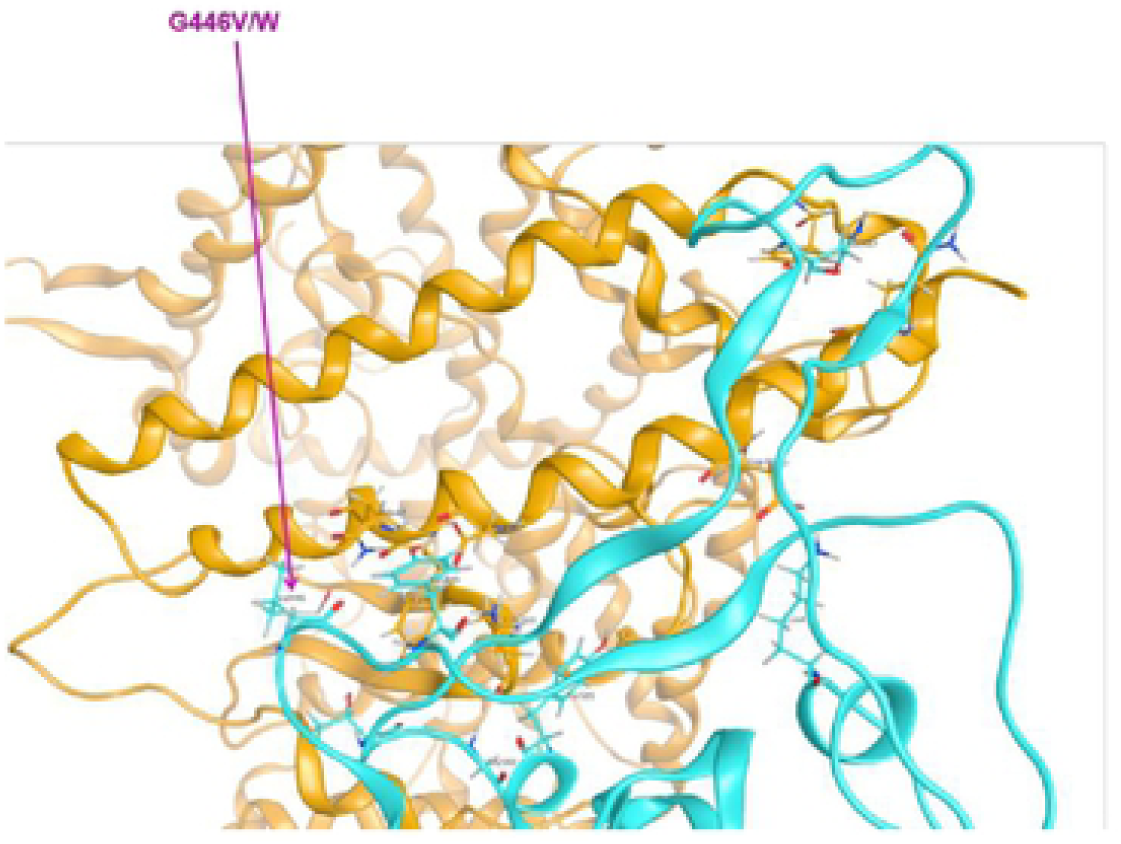
Residue G446 variants in documents from CDC (G446V) and in our prediction (G446W)

Combined 1,150 variants (634 missense mutations in total) in the spike protein of SARS-CoV-2 reported by CDC, we found their affinity and stability in our analysis outcomes. Among 76 missense variants that locate in the region between T333-C525, five variants in spike protein enhance its affinity binding to hACE2 (Table 3) and only three of them increase the stability of spike-hACE2 complex (Table 4). Additionally, we collected the genome sequences of SARS-CoV-2 that harbor those variants in spike protein and found the variants V483A occurs in 26 strains from the USA, V367F occurs in 12 strains from Hong Kong, Australia, and other European countries, and a variant G446V that highly enhance the affinity were identified in a strain from Australia (Supplementary Table 2). We mapped those strains onto our genomic phylogenetic tree and found they were dispersed distributed in the position close to the SARS-bat and SARS-pangolin. As COVID-19 widespread globally, several studies tried to find where this novel coronavirus came from, but few reports analyze the origin of SARS-CoV-2 according to the divergent pattern in spike protein. In our study, the multiple variants in spike protein found in different countries and clustered in the ancestral direction on phylogeny which might suggest SARS-CoV-2 infection start to occur at multiple sites but not only in Wuhan city. However, due to the limited detection capability and restricted availability of samples from infected animals, the variants available in the national database are uncompleted, more data needs to be collected for further studies.

**Table 3.**
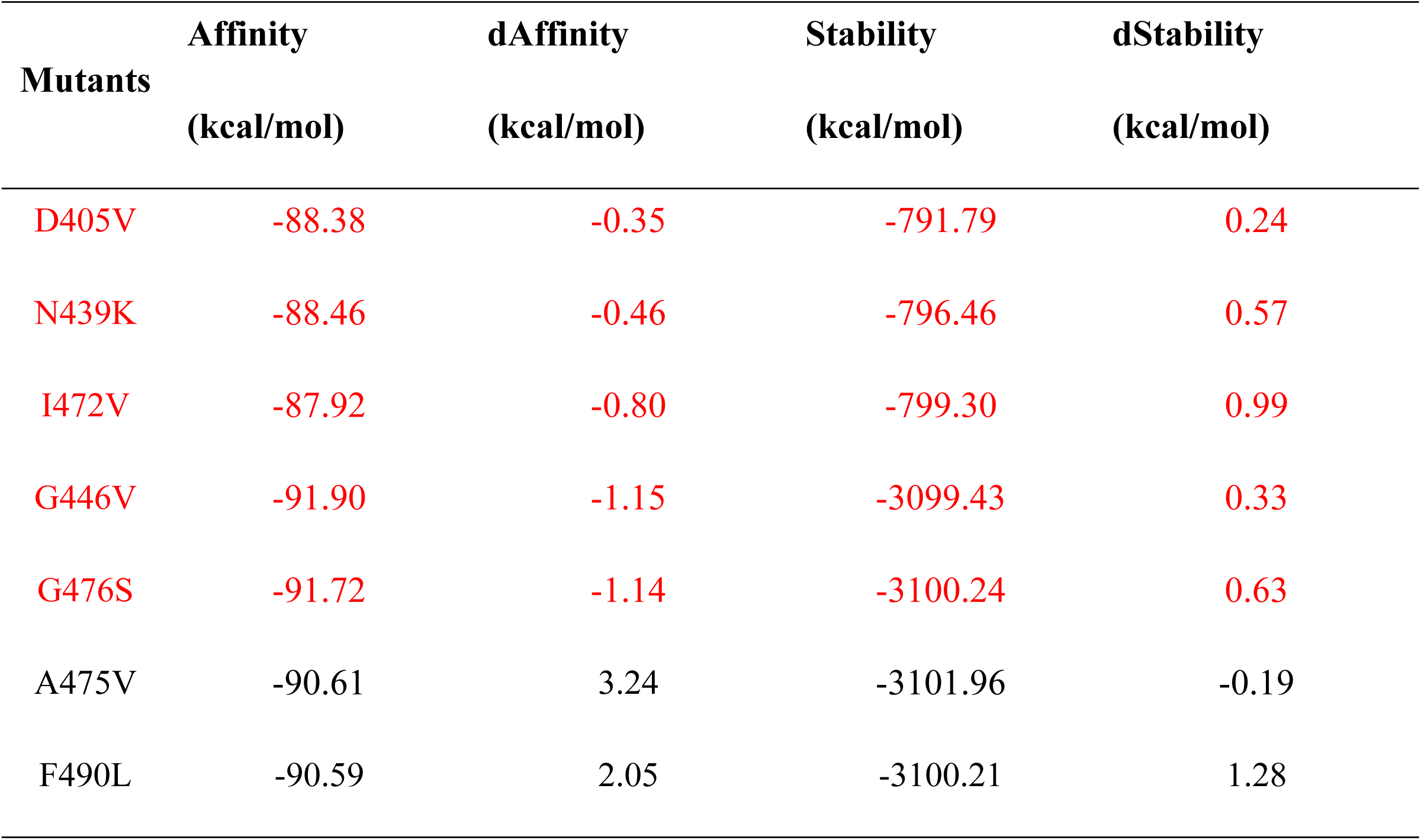
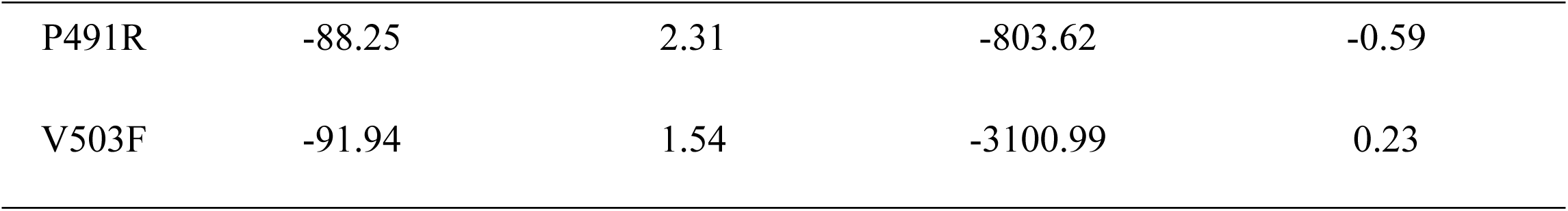
The affinity of the variants of SARS-CoV-2 spike RBD bound to human ACE2, red marks the enhanced ones while black represents the decreased ones.

| Mutants | Affinity<br>(kcal/mol) | dAffinity<br>(kcal/mol) | Stability<br>(kcal/mol) | dStability<br>(kcal/mol) |
| --- | --- | --- | --- | --- |
| D405V | -88.38 | -0.35 | -791.79 | 0.24 |
| N439K | -88.46 | -0.46 | -796.46 | 0.57 |
| I472V | -87.92 | -0.80 | -799.30 | 0.99 |
| G446V | -91.90 | -1.15 | -3099.43 | 0.33 |
| G476S | -91.72 | -1.14 | -3100.24 | 0.63 |
| A475V | -90.61 | 3.24 | -3101.96 | -0.19 |
| F490L | -90.59 | 2.05 | -3100.21 | 1.28 |
| P491R | -88.25 | 2.31 | -803.62 | -0.59 |
| V503F | -91.94 | 1.54 | -3100.99 | 0.23 |

**Table 4.** The stability of the variants of SARS-CoV-2 spike RBD bound to human ACE2, red marks the enhanced ones while black represents the decreased ones.

| **Mutants** | **Affinity**<br>**(kcal/mol)** | **dAffinity**<br>**(kcal/mol)** | **Stability**<br>**(kcal/mol)** | **dStability**<br>**(kcal/mol)** |
| --- | --- | --- | --- | --- |
| D364Y | -88.26 | -0.00 | -792.45 | -0.87 |
| D467V | -88.26 | -0.00 | -793.14 | -0.68 |
| P491R | -88.25 | 2.31 | -803.62 | -0.59 |
| V367F | -88.26 | 6.85E-12 | -799.28 | 1.06 |
| R408I | -88.26 | -0.00 | -792.67 | 1.44 |
| I434K | -88.26 | 1.27E-07 | -800.49 | 1.84 |
| A435S | -88.26 | -4.22E-05 | -799.69 | 1.04 |
| K458N | -88.186 | 0.06 | -793.07 | 1.29 |
| V483A | -88.606 | -0.012 | -797.29 | 1.01 |
| V483I | -88.726 | -0.13 | -797.21 | 1.09 |
| F490L | -90.596 | 2.05 | -3100.21 | 1.28 |
| R509K | -88.26 | 5.45E-05 | -797.38 | 2.04 |
| Y508H | -88.18 | 0.05 | -798.96 | 2.41 |

Furthermore, since the SARS-CoV-2 strains have variants on spike protein relative to a reference sequence of SARS-CoV-2 (NC_045512.2) were found more divergence on the phylogenetic position that closer to SARS-bat and SARS-pangolin, it might indicate a consensus evolution occurred in S gene and enable SARS-CoV-2 to adapt to human hosts well. However, since it is difficult to determine the exact time the zero patient who got infected by SARS-CoV-2, the date of the sample collection and data extracted could mislead the results. We believe further clinical information from all the countries would be needed for research and this is a global concern that requires more cooperation and collaboration but not political games or constant blames. We shared all the analysis of over 3000 variants on spike protein in this study to help the world tracking the mutations of SARS-CoV-2 and also can be useful to select the potential druggable targets and neutral inhibitors to prevent the further damages may be brought by the pandemic of COVID-19.

Aside from mutations in spike protein for adapting to new hosts, the hACE2 also represents polymorphisms in the binding region, host-virus interaction over time makes a natural selection on both virus and host cells ^30^. Therefore, the variants in hACE2 receptor would also play a role in SARS-CoV-2 infection. Cao and his colleagues (2020)^31^ found the polymorphisms of hACE2 do not bring differences in different populations in resisting to the SARS-CoV-2, but their dataset is limited. A study published on bioRxiv shows variants K26R, S16P, T27A, K31R, H34R, E35K, E37K, D38V, N51S, N64K, K68E, F72V, T921, Q102P, G326E, G352V, D355N, H378R, Q388L, and D509Y are predicted to increase the susceptibility of the individuals carrying these variations while K31R, E35K, E37K, D38V, N33I, H34R, Q388L and Y83H decrease binding capacity bind to SARS-CoV-2 spike protein. In our study, we found all the variants in ACE2 from GnomAD and local dataset addition to their population frequency in different groups of human beings, same with Eric’s results^32^, no significant differences in ACE2 variant frequency were found from GnomAD while three high allele frequency hACE2 variants (I468V, N638S and R708Q) were identified in local population with their allele frequencies are very low in other populations (Figure 8). Researchers found three unique variants in hACE2 in the Italian population that might be corresponding to the high fatality including P389H, W69C, and L351V^33^.

**Figure 8.**
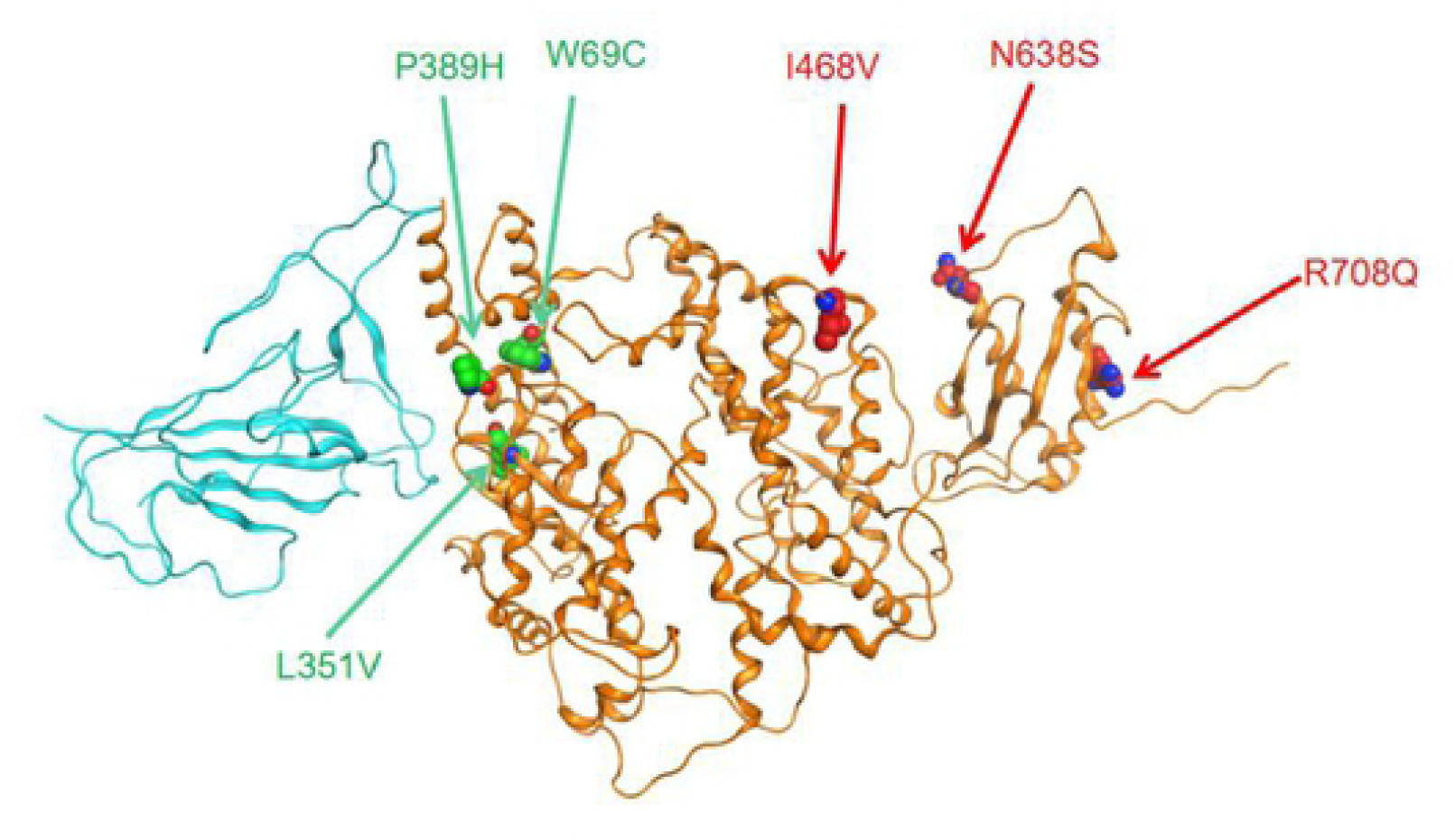
Spike protein polymorphous points from local population and Italy (Citation), the Italian ones are marked as green while local ones marked as red.

We compared the unique variants found in local and Italian population by mapping them on the hACE protein structure (Figure 8) and found the Italian ones were closer to the binding region than local ones, but *in silico* simulation indicated that none of them change the affinity and stability of spike protein of SARS-CoV-2 and hACE2 complex (Table 2). Generally, we did not find any variants in ACE2 that would significantly increase or decrease the affinity and stability of spike protein binding to hACE2 which may indicate SARS-CoV-2 enable to indiscriminately infect all humans. Due to various diet and health conditions in different individuals, the efficiency and virulence of SARS-CoV-2 might be different, and the mortality rate is also more likely to be related to medical and health conditions and the physical condition of patients.

Up to date, there is no effective therapeutics approved universally for both treatment and prevention of COVID-19. It is still urgent to understand the mechanism of the evolution, adaptation and transmission of SARS-CoV-2 and develop an efficient inhibitor or vaccine to prevent SARS-CoV-2 leading to COVID-19. The variants in spike protein of SARS-CoV-2 and hACE2 and the affinity and stability of spike-hACE2 complex would provide a database for tracking the adaptive mutation of SARS-CoV-2 and potential recombination events across different species. A study in review right now shows ORF1ab of SARS-CoV-2 and MERS-CoV may have recombination and result in more severe disease (unpublished). The analysis shared in this study would provide useful genetic information to track the mutations that occur in spike protein of SARS-CoV-2 and prevent the recurrence of this epidemic, moreover, protect human beings from zoonotic coronavirus infection.

## Acknowledgments

We thank the supports provided by the Academy of Medical sciences Zhengzhou University and the data analysis was supported by Supercomputing Center in Zhengzhou University (Zhengzhou). The study is funded by the Collaborative Innovation Project of Zhengzhou (Zhengzhou University) (Grant No. 18XTZX12004) to WT, the Department of Science and Technology of Henan Province (Grants No. 201100312100) to JL, Project of Basic Research Fund of Henan Institute of Medical and Pharmacological Sciences to JS (Grant No.: 2019BP0202). Any opinions, findings, conclusions, or recommendations expressed here are those of the author(s) and do not necessarily reflect the views of the funding agencies.

## Contributions

XX, JXS, HEX designed and conducted the current study. YPQ, ZGY, SSF, DHL provided bio-informatics supports and performed the main analysis. WXT, JCG, YHW, JBL supported the methodological and pathological information. XX, JXS, HEX, YPQ wrote the manuscript. All authors revised and approved the final manuscript.

